# Protection of the human gut microbiome from antibiotics

**DOI:** 10.1101/169813

**Authors:** Jean de Gunzburg, Amine Ghozlane, Annie Ducher, Emmanuelle Le Chatelier, Xavier Duval, Etienne Ruppé, Laurence Armand-Lefevre, Frédérique Sablier-Gallis, Charles Burdet, Loubna Alavoine, Elisabeth Chachaty, Violaine Augustin, Marina Varastet, Florence Levenez, Sean Kennedy, Nicolas Pons, France Mentré, Antoine Andremont

## Abstract

**Background:** Antibiotics are life-saving drugs but severely affect the gut microbiome with short term consequences including diarrhoea, *Clostridium difficile* infections and selection of antibiotic-resistant bacteria. Long-term links to allergy and obesity are also suggested. We devised a product, DAV132, and previously showed its ability to deliver a powerful adsorbent, activated charcoal, in the late ileum of human volunteers.

**Methods:** We performed a randomized controlled trial (ClinicalTrials.gov NCT02176005) in 28 human volunteers treated with a 5-day clinical regimen of the fluoroquinolone antibiotic moxifloxacin in two parallel groups, with or without DAV132 co-administration. Two control goups of 8 volunteers each receiving DAV132 alone, or a non-active substitute, were added.

**Results:** The co-administration of DAV132 decreased free moxifloxacin fecal concentrations by 99%, while plasmatic levels were unaffected. Shotgun quantitative metagenomics showed that the richness and composition of the intestinal microbiota were largely preserved in subjects co-treated with DAV132 in addition to moxifloxacin. No adverse effect was observed. In addition, DAV132 efficiently adsorbed a wide range of clinically relevant antibiotics *ex-vivo*.

**Conclusions:** DAV132 was highly effective to protect the gut microbiome of moxifloxacin - treated healthy volunteers and may constitute a clinical breakthrough by preventing adverse health consequences of a wide range of antibiotic treatments.

## INTRODUCTION

Antibiotics constitute one of the most medically important and effective class of drugs. However, during systemic antibiotic treatments, the non-absorbed part of orally administered drugs, as well as the possible fraction excreted into the upper intestine via bile for both oral and parenteral antibiotics, reaches the cecum and colon where it can exert devasting effects on the gut microbiome, with short and long term consequences [1–5]. Short-term effects include diarrhoea, *Clostridium difficile* infection (CDI) and selection of antibiotic-resistant microorganisms [6–8]. CDI currently constitutes a major clinical challenge, and antibiotic treatments are the key factor for their occurrence in hospitalized and community patients [9–11]. Longterm links to allergy [4] and obesity [5] have also been suggested. Indeed, bourgeonning research in recent years has shown that the intestinal microbiome plays an important role in many aspects of human physiology and health [12]. In particular it is involved in the production of metabolites that may affect insulin sensitivity [13] and diet-related obesity [14]; this state has been shown to correlate with a gut microbiome of lower bacterial richness than in healthy individuals [15].

Strategies that would preserve the intestinal microbiome from deleterious consequences of dysbiosis during antibiotic treatments, would be highly welcome for immediate protection of patients from CDI, and also for long term public health consequences such as dissemination of resistant bacteria and the occurrence of metabolic disorders. Oral administration of a β-lactamase [16–20] prevented the impact of parenteral β-lactams on the microbiome, which is promising but limited to this class of antibiotics. Delivering a non-specific adsorbent to the colon partially decreased fecal concentrations of orally-administered ciprofloxacin without significantly affecting its plasma pharmacokinetics in rats [21]. We devised a product, DAV132, which delivers a powerful non-specific adsorbent, a carefully chosen activated charcoal, to the late ileum in humans, and have shown in healthy volunteers that its administration did not affect the plasma pharmako-kinetics of amoxicillin, given as a single dose [22]. Here, we performed a randomized clinical trial in volunteers receiving a full oral clinical course of the fluoroquinolone antibiotic moxifloxacin (MXF), and assessed DAV132 safety, as well as its resulting effects on MXF plasma concentrations, MXF free fecal concentrations, and intestinal dysbiosis. We also evaluated *ex-vivo* the capacity of DAV132 to adsorb a wide range of clinically relevant antibiotics.

## METHODS

### Investigational products

DAV132 was manufactured according to Da Volterra’s specifications [22] under Good Manufacturing Practice conditions at NextPharma (Bielefeld, Germany). The dosage form, consisting of 7.5 g DAV132, contained 5.11 g activated charcoal as active adsorbing ingredient. To facilitate oral intake, DAV132 pellets were suspended in an extemporaneously prepared gel suspension (Batch number: C1311007). A negative control (CTL) was made of a product similar to DAV132, in which the adsorbent was replaced by microcrystalline cellulose (Batch number: C1311006). MXF was from Bayer (Avalox 400 mg Filmtabletten, batch number: BXGFBN1).

### Subjects and clinical trial design

Male and female healthy volunteers over 18 years old having given written informed consent were included (body mass index < 30 kg/m^2^, normal digestive transit and healthy (by medical history, physical examination, vital signs, electrocardiogram and blood laboratory results) at a screening visit 8-21 days prior to treatment beginning defined as Day 1 (D1)). Subjects carrying *C*. *difficile* at screening or with a history of hospitalization or antibiotic exposure (both past 3 months), or vaccination (past 28 days) were not included.

Volunteers were included as outpatients (March-October 2014) in a prospective, randomized, controlled, repeated doses, open-label trial, blinded to analytical and microbiological evaluations, at the Clinical Investigation Centre of the Bichat Hospital, Paris (France) in respect with Good Clinical Practice and the Declaration of Helsinki as last amended. Volunteers were randomised (Supplementary Material) to receive either MXF alone (n=14), MXF+DAV132 (n=14), DAV132 alone (n=8), or negative control (CTL) (n=8) (Figure 1). The study was carried out after authorisations from French Health Authorities and the Independent Ethics Committee (“Comité de Protection des Personnes Ile-de-France IV”, Paris, France) had been obtained (January and February 2014, respectively). It was declared in ClinicalTrials.gov (identifier NCT02176005) and the French (ID-RCB number 2013-A01504-41) registers of clinical trials. A study-specific Scientific Committee (A.A., V.A., A. Duc., A.Duf., X.D., C.F., J.G., F.M., and M.V.) was set up to ensure scientific integrity.

**Figure 1:**
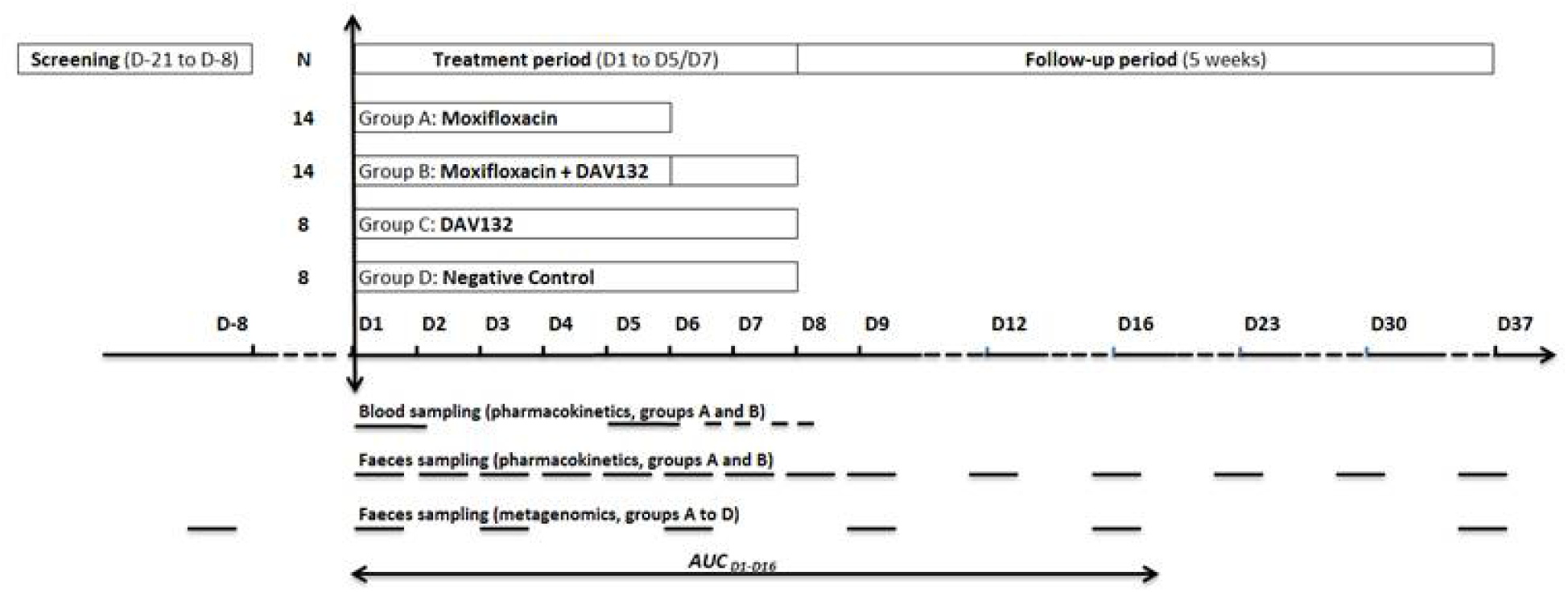
Study design. The various periods of the study (screening, treatment, follow-up) are shown in boxes at the top. The times of blood and feces sampling for MXF pharmacokinetics and metagenomics analysis are shown by horizontal bars in the bottom section of the graph.

### Treatments

MXF 400 mg was administered orally, once a day (od, after breakfast) from D1 to D5 under direct observed therapy. DAV132 7.5 g (or CTL) was administered orally, thrice a day (tid, before meals) from D1 to D7; on D1, the first DAV132 dose was given two hours before MXF. Morning administrations of DAV132 and of CTL were performed under direct observed therapy, while noon and evening intake were reported by the subjects. Compliance was assessed by counting empty bottles each following day. Follow-up was until D37. See Supplementary Material for Collection and storage of fecal and plasma samples.

### MXF assay in plasma and fecal samples

MXF assays were performed by Amatsi Group (Fontenilles, France) using specifically developed and validated bioanalytical methods. MXF concentrations were determined by reversed phase HPLC coupled with fluorescence detection for plasma levels, and tandem mass spectrometry detection for free fecal concentrations (Supplemental Material).

### Statistical methods

The primary objective was to evaluate the influence of DAV132 on free fecal MXF concentrations between D1 and D16 by comparing individuals in the MXF and MXF+DAV132 treated groups. The primary endpoint was the AUC_D1-D16_ of free fecal MXF concentrations. The study sample size was calculated at the time of study design. Assuming an AUC_D1-D16_ variability similar to that of the AUC_D1-D14_ previously measured from individual data [23], a sample size of 11 subjects in each MXF treated group (MXF and MXF+DAV132) would allow to detect a 2-fold change between these groups (90% power, two-sided test, type I error 0.05). For security, we included 14 subjects in each of these groups. Additionally, we randomized two groups of 8 volunteers without MXF, but with DAV132 or CTL to study secondary objectives (DAV132 safety and intestinal microbiota composition).

As preplanned for the primary objective, comparison of log(AUC_D1-D16_) of free MXF fecal concentrations, in groups treated with MXF+DAV132 and MXF alone, was performed using a general linear model. AUC_D1-D16_ were calculated by the trapezoidal method using the actual time of stool emission and the results were expressed as geometric means of AUC_D1-D16_ and coefficient of variation. For MXF plasma concentrations, comparisons of log(AUC_0-24h_) and log(Cmax), in groups treated with MXF+DAV132 and MXF alone, were perfomed using a general linear model. AUC_0-24h_ was calculated by the trapezoidal method. Statistical analysis of clinical and pharmacokinetic data were performed using the SAS 9.4 software (SAS Institute, Cary, NC, USA).

### Metagenomic methods and analysis

Analysis of metagenomic data was exploratory and not prespecified. Essentially, total fecal DNA was extracted as described [24,25] and sequenced using SOLiD 5500 Wildfire (Life Technologies) resulting in 67.2 ±19.8M (mean ±SD) sequences of 35-base-long single-end reads. High-quality reads were generated with quality score cut-off >20. Reads with a positive match with human, plant, cow or SOLiD adapter sequences were removed. Filtered high-quality reads were mapped to the MetaHIT 3.9M gene catalog [26] using METEOR software [27]. The read alignments were performed in colorspace with Bowtie software (version 1.1.0) [28] with options: -v 3 (maximum number of mismatch) and -k 10000 (maximum number of alignment per reads). The raw SOLiD read data were deposited in the EBI European Nucleotide Archive under accession number PRJEB12391. Details of read mapping, data treatment and statistical methods to analyse microbiome data are in Supplementary Material.

### Adsorption of antibiotics by activated charcoal *ex vivo*

To mimic at best the adsorption of antibiotics onto activated charcoal in the gut we used cecal medium that had been obtained from extemporaneously euthanised pigs, and stored at -80°C since. Antibiotics (400 μg/mL), and activated charcoal (4 mg/mL) obtained from DAV132 deformulated by incubation for 30min at 37°C in 50 mM sodium phosphate buffer pH 7.5 containing 80 nM NaCl, were independently pre-incubated with cecal medium (1:1 v:v) for 2h at 37°C. Then, the two pre-incubation reactions were mixed and further incubated for 3h at 37°C with gentle agitation. When the tested antibiotic was sensitive to β-lactamases, endogenous enzymes were inactivated by heating at 70°C for 1h. Samples were centrifuged 3minX19,890 g, and the concentrations of non-adsorbed antibiotics in the supernatant were quantified in triplicate using a microbiological assay [29].

## RESULTS

### Subjects

Overall, 71 subjects were included in the DAV132-CL-1002 study between March 20, 2014 and September 01, 2014. 21 subjects did not meet the inclusion/exclusion criteria and 5 subjects withdrew consent before randomization. Of 45 subjects randomized, one refused to take treatment and withdrew from the study; 44 were treated and completed the study: n=14 in groups MXF and MXF+DAV132, n=8 in groups DAV132 and CTL. All subjects meeting the inclusion/exclusion criteria, having taken at least 20 doses of DAV132 (95% of expected doses) and 5 doses of moxifloxacin (100% of expected doses) were evaluable and included in the per protocol population. There were no deviations from protocol during the treatment and follow-up periods. Therefore, the 44 subjects were included in both per protocol and safety analysis sets. The number of subjects analyzed in groups MFX and MFX+DAV132 ensured a study statistical power over 90%; the characteristics of volunteers were similar in both groups (Supplementary Table 2).

### MXF pharmacokinetics in feces and plasma

In volunteers with MXF alone, average fecal concentrations of free MXF peaked at 136.2 μg/g (with 39% intersubject coefficient of variation, CV) at day 6 (D6) and returned to undetectable levels by D16 (Figure 2a); they were markedly reduced in volunteers that received MXF together with DAV132, with free fecal MXF concentrations ranging from 1 to 14 μg/g faeces between D1 and D6. Indeed, co-administration of DAV132 reduced the AUC_D1-D16_ of fecal free MXF by more than 99%, with geometric means of 699.2 μg/g.day (CV 41%) in the MXF group vs. 6.4 μg/g.day (CV 69%) in the MXF+DAV132 group (p=3.10^−18^). Despite the low concentrations of free fecal MXF in volunteers that were co-administered DAV132, no selection for resistance in coliforms was seen; some quinolone- and fluoroquinolone-resistant strains emerged, but no difference was observed between the treatment groups (Supplementary Table 1). When adjusting for each main individual characteristic of the volunteers (Supplementary Table 2) in a multivariate analysis, the effect of DAV132 on reducing logAUC_D1-D16_ of free fecal MXF concentration remained significant (analysis not shown). By contrast, plasma concentrations of MXF at D1 and D5 were not significantly different in volunteers that received DAV132 or not, in addition to MXF, as shown by analysis of the geometric means of the AUC_0-24_ and Cmax (Figure 2b, c and Table 1).

**Figure 2.**
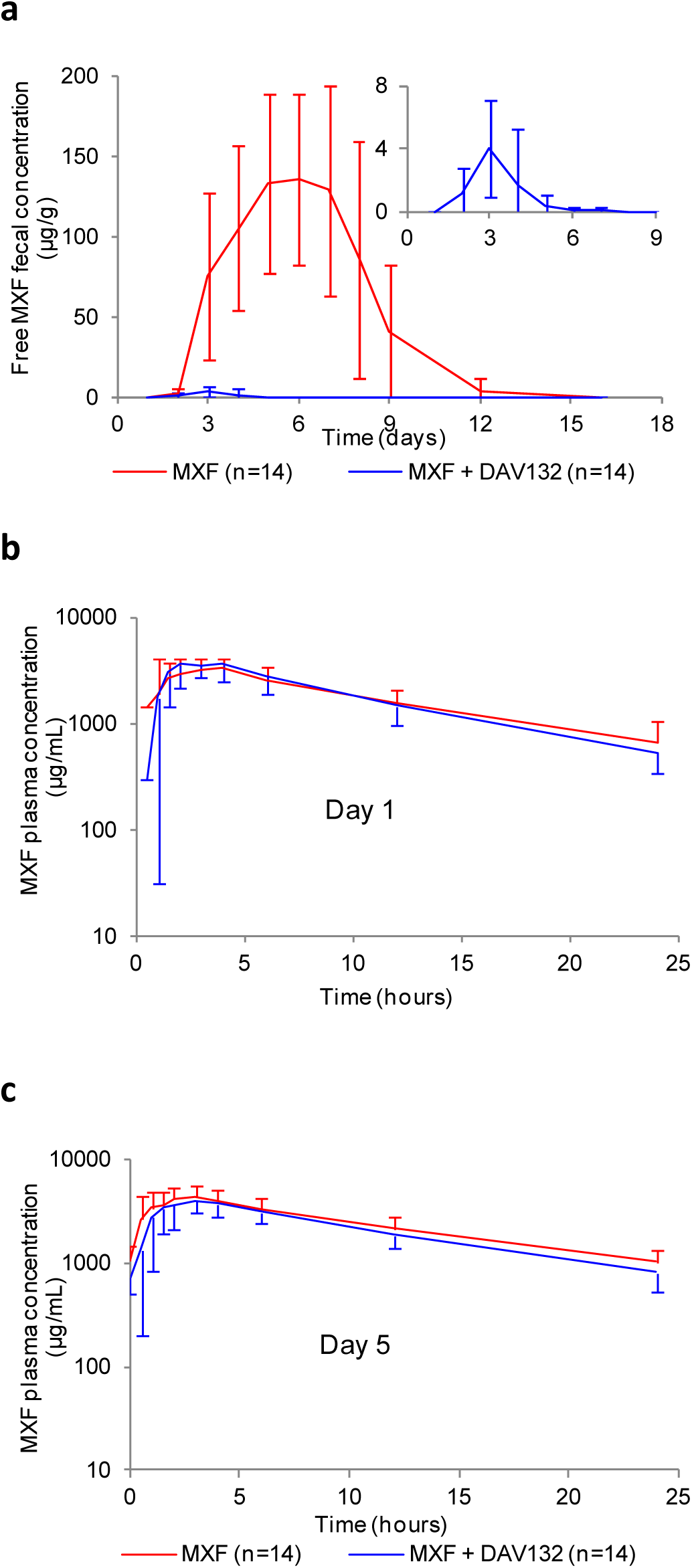
Effect of DAV132 on MXF concentrations in feces and plasma of human volunteers. **(a)** Free fecal MXF concentrations between D1 and D16 (p=10^−17^ for the comparison of logAUC_D1-D16_). Inset: magnified scale for HVs treated with MXF+DAV132. **(b)** Plasma MXF concentrations on D1 (p=0.8 for the comparison of logAUC_0-24h_) and **(c)** D5 (p=0.1 for the comparison of logAUC_0-24h_). HVs, 14 in each of these groups, were administered orally MXF 400 mg od from D1 to D5 (MXF), or MXF 400 mg od plus DAV132 7.5 g tid from D1 to D5 and then DAV132 alone on D6-D7 (MXF+DAV132). Mean values ± standard deviation (SD) are shown.

**Table 1.**
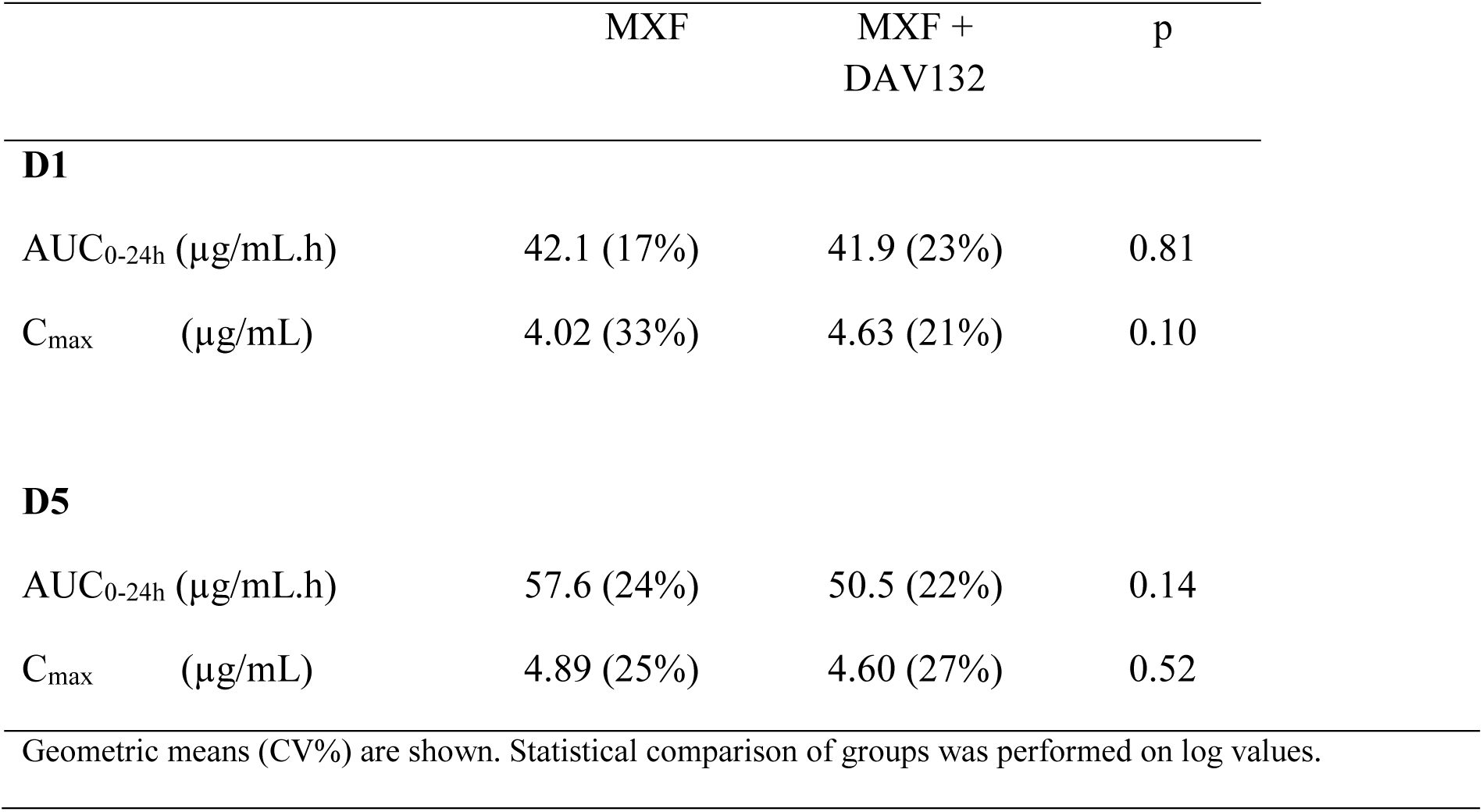
Plasma pharmacokinetic parameters of MXF in HVs receiving or not DAV132.

|  | MXF | MXF +<br>DAV132 | p |
| --- | --- | --- | --- |
| **D1** |  |  |  |
| AUC<sub>0-24h</sub> (µg/mL.h) | 42.1 (17%) | 41.9 (23%) | 0.81 |
| C<sub>max</sub> (µg/mL) | 4.02 (33%) | 4.63 (21%) | 0.10 |
| **D5** |  |  |  |
| AUC<sub>0-24h</sub> (µg/mL.h) | 57.6 (24%) | 50.5 (22%) | 0.14 |
| C<sub>max</sub> (µg/mL) | 4.89 (25%) | 4.60 (27%) | 0.52 |
| Geometric means (CV%) are shown. Statistical comparison of groups was performed on log values. |  |  |  |

The safety analysis showed that repeated oral administration of DAV132 during 7 days was safe and well tolerated. Only one adverse effect, a per-treatment vulvovaginal mycotic infection, possibly related to MXF, was considered as related to a study product by the investigator. No adverse effect considered as related to DAV132 was reported. No clinically relevant abnormality in vital signs, 12-lead ECG parameters and laboratory results occurred in any subject during the study.

### Prevention of intestinal dysbiosis by DAV132

The global effect of DAV132 administration on the gut microbiome was explored in two ways, by assessing microbiome bacterial gene richness and overall composition. Richness was strongly decreased to 54.6% of baseline value at D6 in volunteers who received MXF alone, and failed to return to the initial value even at D37 (Figure 3a); this decrease was greatly attenuated by co-administration of DAV132 (97.8% of baseline value at D6, close to what was observed for the CTL group). We also assessed the impact of treatments on bacterial gene richness over the length of the trial by computing the AUC, between D0 and D16, of its relative change from D0 for each individual (Figure 4a). This interval was chosen because most of richness evolution took place within it, and no residual antibiotic was present at D16 (Figure 2a). The AUC_D0-D16_ of gene richness change was significantly different among the four groups of volunteers (p=4.10^−6^) (Figure 4a). It was significantly lower in volunteers receiving MXF alone than in those in the CTL group (q=1.10^−5^); it was significantly higher in those treated with MXF+DAV132 than in those receiving MXF alone (q=4.10^−7^), and not significantly different from those in the CTL group (q=0.8), thereby showing the protective effect of DAV132 (Figure 4a). Finally, the AUC_D0-D16_ of gene richness change was highly correlated with the AUC_D1-D16_ of free MXF fecal concentrations (Figure 4b), further illustrating the impact of MXF on richness.

**Figure 3:**
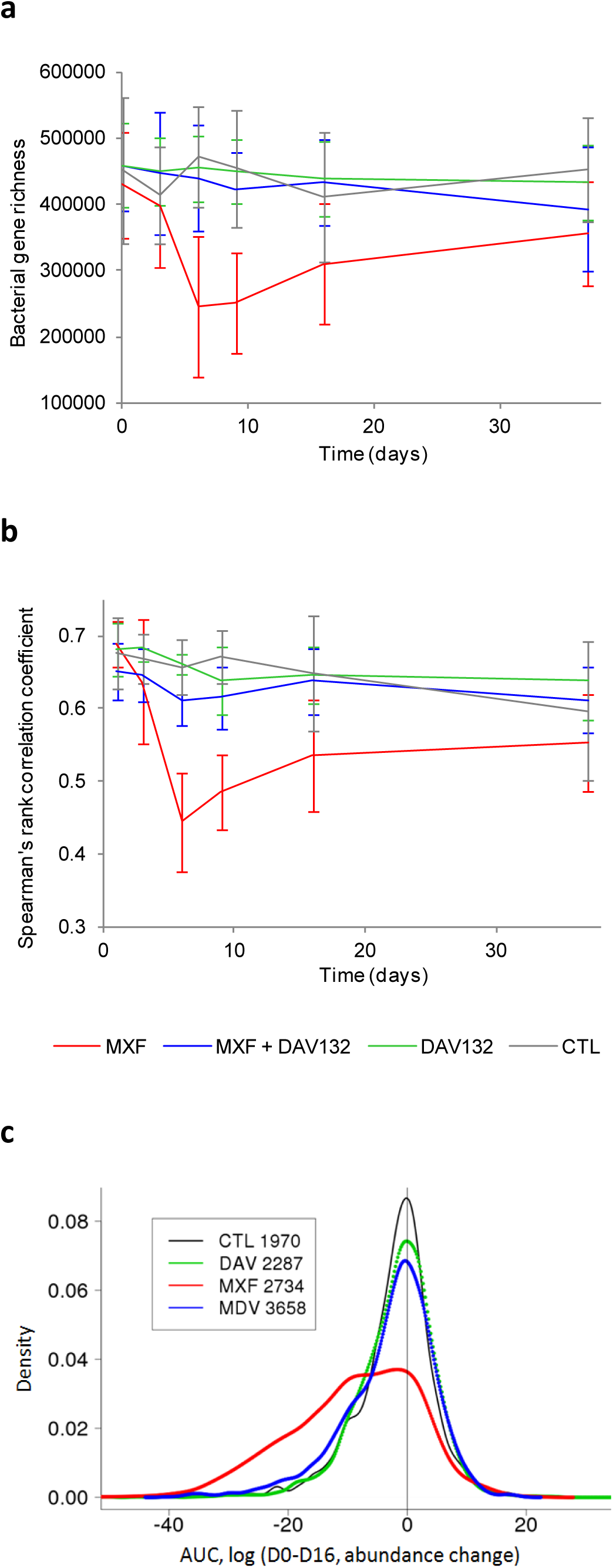
Effect of DAV132 on MXF-induced alterations of the human gut microbiome of human volunteers. **(a)** Gene richness: bacterial gene counts for each study group are displayed. **(b)** Microbiome composition. Spearman’s rank correlation coefficients (ρ) computed from the abundance of all genes of the 3.9 M gene catalog carried by each individual between Dscreening and the indicated days are shown for each study group. **(c)** Metagenomic species. Distribution of AUC values computed between D0 and D16 using log10 of abundance change from D0 of each MGS present in each individual is shown. The group sizes were: MXF n=14; MXF+DAV132 n=13; DAV132 n=8; CTL n=8. The number of available individual measures over all MGS for each group is indicated in the inset. Red, blue, green and black correspond to MXF, MXF+DAV132 (MDV), DAV132 (DAV) and CTL groups, respectively.

**Figure 4:**
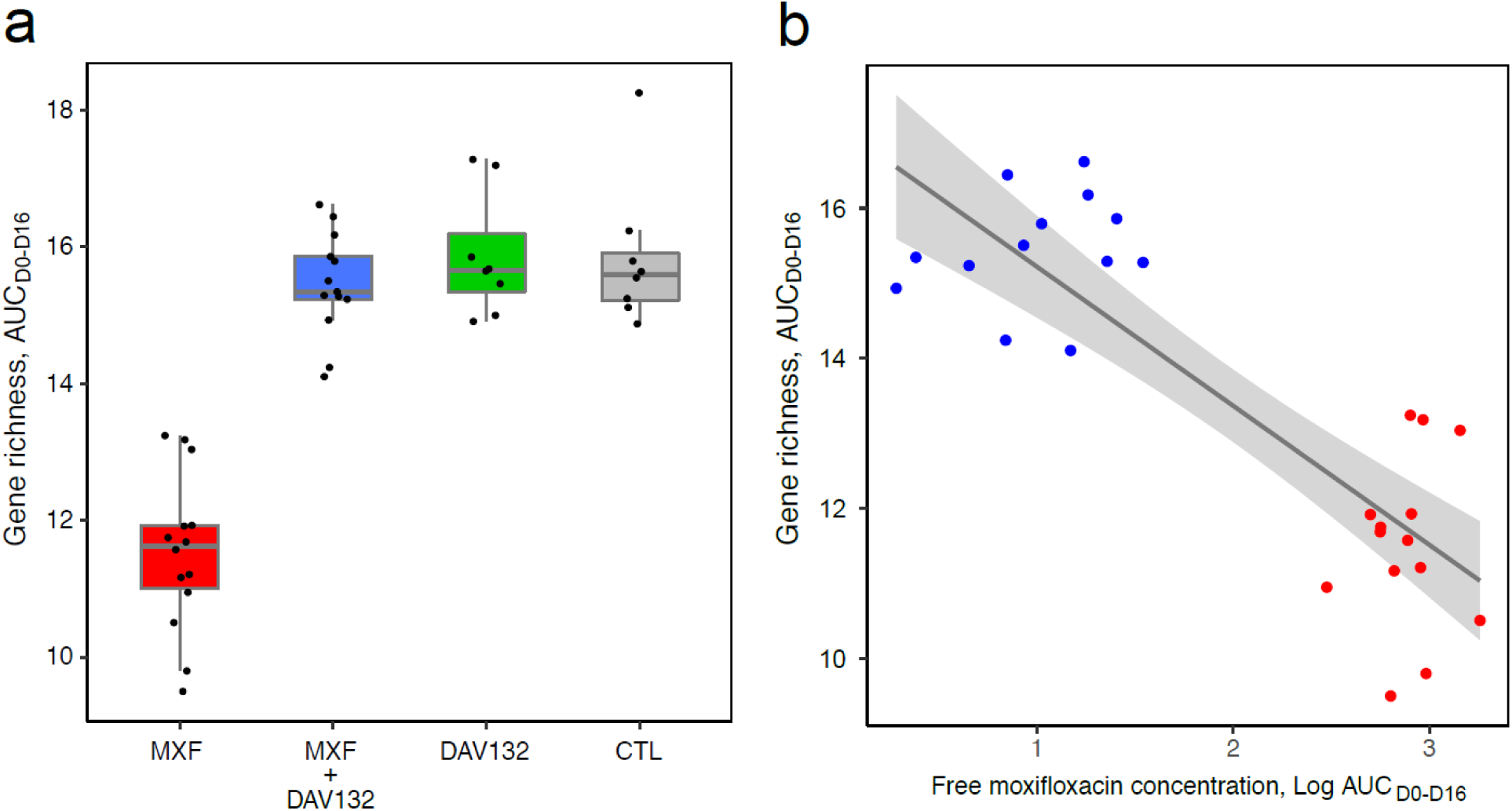
Impact of MXF and DAV132 on gene richness. **(a)**AUC_DO-DI6_ of gene richness change from D0; see methods for details. Medians [min ; max] were 11.63 [9.50 ; 13.24] for MFX, 15.34 [14.10 ; 16.62] for MXF+DAV132, 15.66 [14.91 ; 17.28] for DAV132 and 15.59 [14.88 ; 18.25] for CTL. Of note, in absence of any change from D0 the value of AUC_DO-D16_ would be 16. Median values, quartiles, and 1.5 interquartile range are shown. The distribution of the AUC_DO-D16_ of gene richness was significantly different between the 4 groups (Kruskal-Wallis test p=4.10^−6^). In the pairwise comparisons, it was significantly lower in HVs receiving MXF alone than in those receiving MXF+DAV132 (q=4.10^−7^) or negative control (q=1.10^−5^), whereas the difference between HVs receiving MXF+DAV132 and negative control was not significant (q=0.8) as assessed by the Wilcoxon rank sum test with Benjamini-Hochberg correction for the four pairwise comparisons. No difference was observed between the group receiving DAV132 alone and CTL (q=0.8). The number of individuals in different groups was MXF n=14; MXF+DAV132 n=13; DAV n=8; CTL n=8. **(b)**Relationship between AUC_DO-D16_ of gene richness change from D0 and AUC_D1-D16_ MXF fecal concentration (R^2^=0.71, p=4.10^−8^). Red and blue dots correspond to groups exposed to MXF or MXF+DAV132, respectively.

Overall changes of microbiome composition with time were assessed by computing, for each individual, the Spearman’s rank correlation coefficient of the relative abundance of bacterial genes between each time point and the screening pre-treatment day (Figure 3b). Microbiome composition changed little over time in the CTL and DAV132 groups of volunteers that did not receive MXF (Figure 3b). By contrast, exposure to MXF resulted in marked microbiome changes, detected from D3, maximal at D6, and still partially present 30 days after the treatment ended; these changes were greatly attenuated by DAV132 (Figure 3b). The comparison of Spearman’s rank correlation coefficient values at D6 across the four groups was significant (p=2.10^−6^). Those values were significantly lower in the MXF-treated group (median [min ; max] 0.44 [0.34 ; 0.59]) than in the CTL group (0.65 [0.60 ; 0.70], q=2.10^−4^) and in the MXF+DAV132 group (0.62 [0.53 ; 0.66], q=1.10^−5^). DAV132 exerted an important, but not total protection of the microbiome from the effect of MXF, as the median in the MXF+DAV132 group was slightly lower than in the CTL group (q=0.03).

### Intestinal microbiota analysis at the species level

629/741 (85%) metagenomic species (MGS, see Supplementary Materials) found in the MetaHit gut microbial catalogue of 3.9 M genes were present in at least one sample. The mode of their AUC_D0-D16_ distribution was 0 in the CTL group as well as in volunteers treated with DAV132 alone, indicating that the abundance of most MGS did not change (Figure 3c). The distribution of AUC values for the MXF group was strikingly different, with a broad shoulder toward negative values, indicating a decrease in the abundance of numerous MGS. This shoulder was largely absent in the MXF+DAV132 group, suggesting that many MGS were protected by DAV132.

A detailed analysis of the MGS that differed significantly between treatment groups ( Supplementary Material) showed that of the 252 MGS present at baseline in at least 4 volunteers per group, 99 were differentially abundant. Only 3 were affected by DAV132 given alone, whereas 86 were affected by the MXF treatment; among them, 81% were fully protected, and a further 12% partially protected from the effect of MXF by co-administration of DAV132.

### Taxonomical analysis

Taxonomical characterisation at the genus level (Figure 5) showed that *Alistipes*, *Bilophila*, *Butyciromonas*, *Coprobacillus*, *Fecalibacterium*, *Odoribacter*, *Oscillibacter*, *Parasutterella*, *Roseburia*, and *Sutterella* genera were decreased in MXF-treated volunteers, and partially *(Bilophila*) or fully (all others) protected by DAV132. In contrast, *Bacteroides*, *Paraprevotella* and *Lachnoclostridium* were unaffected by MXF as well as MXF+DAV132 treatments. We used a set of 34 MGS characteristic of the high richness microbiome of healthy individuals present among the 252 examined above (Supplementary Table 4) to assess whether MXF may induce not only an overall loss of gut microbiome richness but also a shift to a composition expected in low richness microbiomes. The AUC_D0-D16_ of log10 of relative abundance change from D0 of these 34 MGS was significantly different among the 4 groups of volunteers (p<10^−4^; Supplementary Figure 2), being significantly lower under MXF treatment alone than under CTL (q=1.10^−4^) or MXF+DAV132 (q=1.10^−4^); just as for overall gene richness, this measure also showed no significant difference between MXF+DAV132 and CTL groups (q=0.6), further attesting of the protective effect of DAV132.

**Figure 5:**
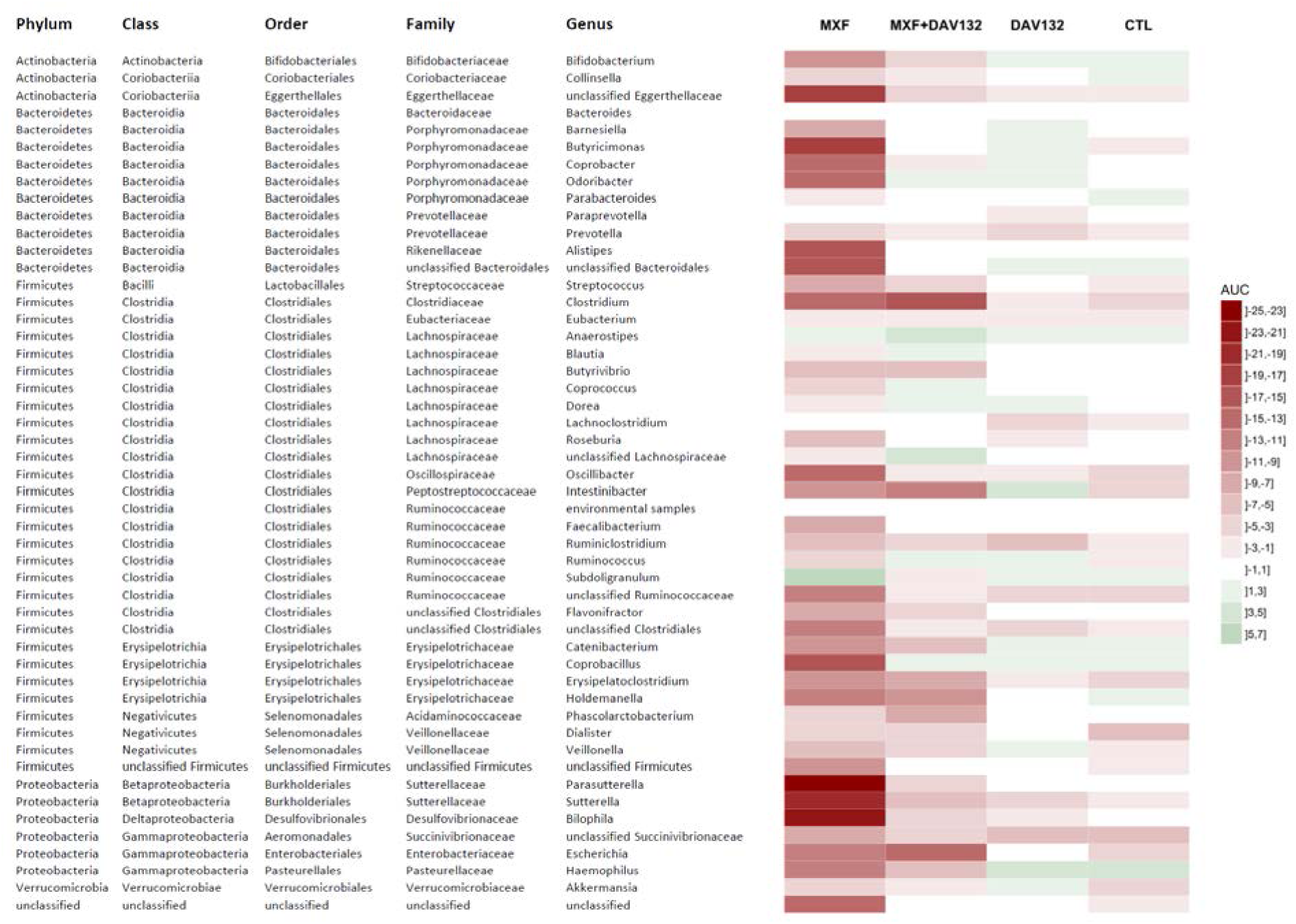
Heatmap depicting the changes induced by the various treatments at the genus level. The heatmap represents, for each treatment group and for each bacterial genus, the median AUC of log10 change from D0 of the MGS that constitute this genus. Green and red coulours respectively indicate genus that are decreased or increased (the intensity of the colour represents the extent of the change), while white indicates very limited changes.

### *Ex-vivo* adsoption of other antibiotics

To assess whether DAV132 could also protect against antibiotics other than MXF routinely used in the clinic, we examined the capacity of the activated charcoal released from DAV132 to adsorb them under *ex vivo* conditions, i.e. in pig cecal medium (Table 2). Among the 14 antibiotics routinely used in clinic tested, 13 were adsorbed to an extent of at least 95% by the charcoal after 3-5h of contact with deformulated DAV132. Only amoxillin was a little less adsorbed, to the extent of 92%.

**Table 2.** Adsorption of antibiotics by deformulated DAV132 *ex vivo*.

| Antibiotic class | Antibiotic | Adsorption by deformed DAV132 at 3 hours and 100:1 DAV132:antibiotic ratio (%) |
| --- | --- | --- |
| Penicillins | Amoxicillin | 92.4* |
|  | Piperacillin | 95.4* |
| Cephalosporins, 1<sup>st</sup> generation | Cefalexine | 97.6 |
| Cephalosporins, 3<sup>rd</sup> generation | Cefotaxime | 96.2* |
|  | Ceftriaxone | 99.4 |
| Carbapenems | Ertapenem | 98.0 |
|  | Imipenem | 99.7 |
|  | Meropenem | 98.1** |
| Fluoroquinolones | Ciprofloxacin | >99.9 |
|  | Levofloxacin | >99.9 |
|  | Lomefloxacin | >99.4 |
|  | Marbofloxacin | >99.9 |
|  | Moxifloxacin | >99.7 |
| Lincosamides | Clindamycin | >99.4 |

\* Adsorption at 5 hours
\*\*Adsorption at 2 hours

## DISCUSSION

Our most important result was that in human volunteers treated with a clinical five-day course of oral MXF, DAV132 spared the intestinal microbiome from exposure to free MXF by over 99%, without affecting the plasma pharmacokinetics of the antibiotic or causing any serious adverse effects. This is a major advance over what we showed in the first DAV132 phase 1 trial that was limited to a small number of volunteers treated with DAV132 for 24h and receiving only a single dose of amoxicillin [22]. Here DAV132 was associated with a full 5 day clinical course of a widely used fluoroquinolone antibiotic. The facts that all randomized volunteers completed the study, and only small amount of data were missing ensure the validity of the results. We conclude that the co-administration of DAV132 with MXF is safe, and should not affect the systemic therapeutic effects of oral, as well as parenteral antibiotic treatments. The fact that DAV132 is able to markedly reduce fecal MXF concentrations without significantly affecting systemic exposure to the antibiotic, contrarily to the use of non-formulated activated charcoal [30], is due to the targeted delivery of the adsorbent component to the ileo-cecal region [22].

Our second most important result was that co-administration of DAV132 largely protected richness and composition of the intestinal microbiota of MXF-treated volunteers. The changes observed with MXF alone that were maximal after 6 days of antibiotic, and persisted a month after the treatment ended were reminiscent of those previously observed with ciprofloxacin [2]. Under co-administration of DAV132 with MXF, they were largely reduced and return to baseline was observed 11 days after treatment ended suggesting that long term consequences of antibiotics might be spared. Indeed, a third of the 252 MGS identified in the gut microbiota of the volunteers were affected by MXF, but the co-administration of DAV132 fully protected 81% of the affected MGS, and a further 12% partially. Of particular interest in that respect was that the 34 MGS that had previously been shown to be associated with the high richness microbiome in healthy individuals [15] were well protected.

A third important result from the study was that the adsorbent released from DAV132 could efficiently adsorb under *ex vivo* conditions mimicking the cecum, antibiotics from several distinct and therapeutically important classes such as β-lactams of all categories (penicillins, cephalosporins and carbapenems), fluoroquinolones and lincosamides. This indeed suggests that the co-administration of DAV132 could protect the human gut microbiome against the deleterious effects of many antibiotics, including those administered orally, without affecting their plasma pharmacokinetics, as we previously showed with the β-lactam amoxicillin [22], and here with the fluoroquinolone MXF. The non-specific nature of the adsorbent used in DAV132 might indeed be advantageous over the use of the recently proposed β-lactamase for prevention of intestinal dysbiosis and *Clostridium difficile* infections which is limited to association with parenteral treatments by penicillins and cephalosporins [20].

Another set of results in this trial was strongly favorable for the possibility to further use DAV132 in the clinic. First concerning safety, in spite of the relatively important dose of charcoal that was given (7.5g tid), the treatment was associated with no significant side effects, in particular intestinal ones but for the black darkening of feces. DAV132 had no impact on blood electrolytes or coagulation parameters, suggesting that it did not interfere with electrolytes exchanges or vitamin K production, which take place in the colon. Compliance to treatment was not an issue for the volunteers. Second, we did not observe any remarquable differential modification in the emergence of quinolone/fluoroquinolone resistant coliforms between the groups of volunteers, even when the free fecal antibiotic concentrations were low as in those that received MXF+DAV132. Notwithstanding the small number of subjects involved in this study, this is reassuring because some have suggested that low concentrations of fluoroquinolones might be prone to increase the selection of resistant bacteria [31].

In spite of these positive results, our work has limitations. First, in this phase 1 trial we did not address directly the efficacy of DAV132 to protect patients againt immediate consequences of antibiotic treatments such as *C*. *difficile* colitis. However, we recently published pre-clinical results in hamsters that suggest that such might well be the case [31]. Second we did not address the possibility that DAV132 might interfere with other drugs that could be taken concomitantly for therapeutic purposes by patients treated with antibiotics. This was far beyond the purpose of the current study but will have to be determined before testing the product in actual patients.

Whatever these limitations, the results of this phase 1 trial appear promising: DAV132 may constitute a breakthrough product to prevent short- and long-term detrimental effects of antibiotic treatments. Further studies are under way to validate the potential of DAV132 in a clinical set-up.

## Funding

The randomised clinical trial was sponsored by Da Volterra (Paris) and funded in part by the European Union Seventh Framework Programme (FP7-HEALTH-2011-single-stage) under grant agreement n° 282004, EvoTAR. Additional funding was from the Metagenopolis grant ANR-11-DPBS-0001, and from BPIFrance under the NOSOBIO collaborative programme.

## Acknowledgements

We thank M. Ghidi and M.N. Bouverne (Da Volterra), as well as C. Féger (Emibiothech) for coordination and management of the clinical study, E. Arcaraz, E. Desmartin, A. Toutin, T. Mezzasalma and C. Toutin (Amatsi Group) for the development and validation of bioanalytical methods, and for performing the fecal and plasma pharmacokinetic analyses, I. Wieder and C. Bourseau for performing the phenotypic microbiological analyses, Pr A. Dufour for biopharmacological analysis, P. Clerson (Orgamétrie) for statistical analysis of the clinical data, N. Galleron and B. Quinquis (Metagenopolis) for generating sequencing data and P. Leonard (Metagenopolis) for informatics support.

## Notes

**Potential conflicts of interest:** AD, FSG, VA and MV are employees of Da Volterra. JG, ER, CB, EC, FM and AA are consultants for Da Volterra. JG, AD, FSG and VA are shareholders of Da Volterra. All authors have submitted the ICMJE Form for Disclosure of Potential Conflicts of Interest.

